# IFN-inducible Human Phospholipid Scramblase 1 (PLSCR1) Protein Restricts HIV-1 Infection by Inhibiting Membrane Fusion

**DOI:** 10.1101/2025.06.23.661195

**Authors:** Yajie Liu, Pei Li, Yi-Min Zheng, Yuta Hikichi, Sherimay Ablan, Eric O. Freed, Shan-Lu Liu

## Abstract

Human phospholipid scramblase 1 (PLSCR1) is an interferon-stimulated gene (ISG) that inhibits viral infections through various mechanisms. Here, we identify PLSCR1 as a host restriction factor that inhibits HIV-1 entry by impairing membrane fusion mediated by the envelope glycoprotein (Env). Using multiple cell types including the human SupT1 T cell line and purified CD4^+^ T cells, we demonstrate that PLSCR1 inhibits the replication of HIV-1 with diverse tropisms and subtypes, as well as HIV-2 and SIV. Mechanistically, we find that PLSCR1 blocks viral entry and cell-to-cell transmission by restricting HIV-1 virion-cell and cell-cell fusion without affecting CD4 or CXCR4 expression or virus binding to the cell surface. Notably, PLSCR1-mediated restriction of viral entry is independent of type I interferon signaling. Collectively, these findings establish PLSCR1 as a broad-spectrum lentiviral restriction factor that acts at the membrane fusion stage, thereby expanding our understanding of ISG-mediated antiviral defense.

## INTRODUCTION

Interferons (IFNs) stimulate broad antiviral responses by inducing a network of IFN-stimulated genes (ISGs), many of which restrict viruses at specific stages of the replication cycle. Phospholipid scramblase 1 (PLSCR1) is an ISG induced by both type I and type II interferons, and has been shown to restrict a range of DNA and RNA viruses, including HBV, EBV, HCMV, HTLV-1, IAV and SARS-CoV-2 (1-6). PLSCR1 is a type II membrane protein anchored at the plasma membrane via its C-terminal tail and possesses a scramblase activity (7). Many of its antiviral activities have been linked to interactions with viral proteins, such as HBV HBx, EBV BZLF1, CMV IE2, or HTLV-1 Tax (1, 3, 5, 6, 8). Recently, PLSCR1 has been reported to restrict SARS-CoV-2 by targeting spike-containing vesicles and preventing membrane fusion on the plasma membrane and/or endosomal compartments (4, 9). While prior studies have reported interactions between PLSCR1 and the HIV-1 Tat protein or cellular CD4 (10, 11), whether or not HIV replication is restricted is currently unknown.

Infection of HIV and related lentiviruses is limited by numerous host restriction factors, including APOBEC3G, TRIM5α, Tetherin, SAMHD1, IFITM, MxB, SERINC and TIM (12-15). Herein, we investigated the role of PLSCR1 in HIV-1 infection in various cells, including the SupT1 T cell line and purified CD4^+^ T cells. We found that PLSCR1 inhibits HIV-1 entry and cell-to-cell transmission by blocking membrane fusion, without affecting CD4/CXCR4 expression or viral binding to target cells. This restriction extends to other lentiviruses, including primary HIV-1 isolates, HIV-2, and SIV.

## RESULTS AND DISCUSSION

We generated SupT1 stable cell lines using lentiviral vectors expressing shRNA control or targeting human PLSCR1. Cells were infected with 5 ng p24 of HIV-1 NL4-3, and viral replication was monitored by measuring the levels of p24 in newly produced virions every 2 days over a 14-day period. PLSCR1 knockdown (KD) enhanced HIV-1 replication, as evidenced by elevated p24 levels that peaked on day 8, compared to shRNA control cells where p24 levels remained relatively low and peaked on day 12 (**Fig. 1A**). We also treated SupT1 cells expressing either control or PLSCR1-targeting shRNA with 0, 50, or 200 U/ml of IFN-β prior to HIV-1 infection. Western blot analysis confirmed that PLSCR1 expression was upregulated by IFN-β, and its antiviral activity was maintained, as evidenced by increased Gag expression in PLSCR1-depleted cells (**Fig. 1B**). These results demonstrate that both constitutively expressed and IFN-induced PLSCR1 intrinsically restrict HIV-1 replication.

**Figure 1.**
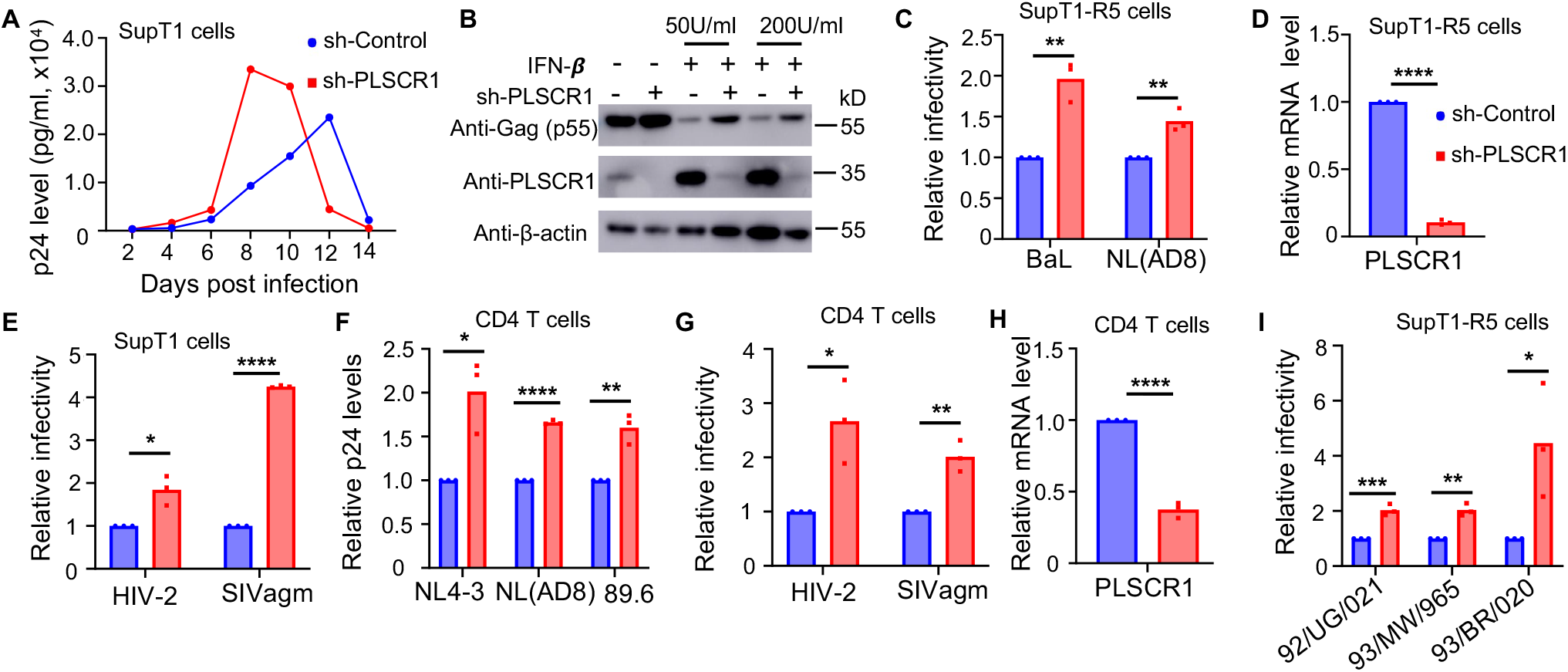
PLSCR1 restricts the replication of HIV-1, HIV-2, and SIV. **(A)** Replication kinetics of HIV-1 NL4-3 in SupT1 cells stably expressing shControl or shPLSCR1 as measured by p24 ELISA. **(B)** Western blotting analysis showing induction of PLSCR1 expression by IFN-β and increased Gag levels by shPLSCR1 knockdown in SupT1 cells compared to controls. **(C)** Relative infectivity of HIV-1 BaL and AD8 produced in SupT1-R5 cells transiently transduced with lentiviral vector expressing shControl or shPLSCR1. **(D)** The mRNA knockdown efficiency of PLSCR1 in SupT1-R5 cells determined by qPCR. **(E)** Relative infectivity of HIV-2 and SIVagm in HeLa-TZM cells. **(F)** Replication of HIV-1 NL4-3, AD8, and 89.6 in CD4^+^ T cells (three donors, n=3) examined by p24 ELISA. **(G)** Relative infectivity of HIV-2 and SIVagm in primary CD4^+^ T cells. (**H**) The PLSCR1 mRNA knockdown efficiency in primary CD4^+^ T cells. **(I)** Relative infectivity of three primary HIV-1 subtypes in SupT1-R5 cells with or without PLSCR1 knockdown. Data represent means ± SEM from three or more independent experiments. Statistical significance was determined using unpaired two-tailed Student’s t-test. *p < 0.05, **p < 0.01, ***p < 0.001, ****p < 0.0001.

To determine whether this restriction extends to CCR5-tropic (R5-tropic) strains of HIV-1, we transiently transduced SupT1-R5 cells with PLSCR1-targeting shRNA and infected them with R5-tropic proviral HIV-1 isolates NL(AD8) and BaL for 2 days. We observed that replication by BaL and NL(AD8) was increased by 2.0-fold (p < 0.01) and 1.5-fold (p < 0.01), respectively, in PLSCR1 KD cells compared to shRNA control cells (**Fig. 1C-D**). Similar restriction by PLSCR1 was observed for HIV-2 and the primate lentivirus SIVagm in SupT1 cells, which exhibited ∼2-fold (p < 0.05) and ∼4-fold (p < 0.001) increased infectivity, respectively (**Fig.1E**). In addition, we validated the restriction phenotype in purified human CD4^+^ cells for HIV-1 X4-tropic NL4-3, R5-tropic NL(AD8), and dual-tropic 89.6 (**Fig.1F**), as well as HIV-2 and SIVagm (**Fig.1G**). The PLSCR1 knockdown efficiency was robust in CD4+ cells (**Fig. 1H)**. We extended our analysis to primary isolates representing different HIV-1 subtypes and found that PLSCR1 inhibited replication of subtype A 92/UG/021 (X4-tropic), subtype C 93/MW/965 (R5-tropic), and subtype F 93/BR/020 (dual-tropic, R5/X4), with the strongest restriction observed against 93/BR/020 (**Fig. 1I**). Collectively, these results demonstrate that PLSCR1 broadly restricts lentivirus replication.

To determine if PLSCR1 acts on the entry of HIV-1, we applied HIV-1 pseudovirions encoding GFP (pLenti-puro-GFP) and bearing different HIV-1 Env, i.e., NL4-3, BH10, or HXB2. We found that knockdown of PLSCR1 in SupT1 cells increased HIV-1 entry mediated by all three Envs (**Fig. 2A**), indicating that PLSCR1 inhibits HIV-1 entry. We next examined whether PLSCR1 affects HIV-1 virion binding to the target cell surface by incubating the purified HIV-1 Gag-GFP virus-like particles (VLPs) bearing NL4-3 Env with SupT1 shRNA-Control or shPLSCR1 cells on ice for 2 hours, followed by determining cell-associated GFP-virion signals using flow cytometry after extensive washes. As shown in **Fig. 2B**, PLSCR1 knockdown had no measurable effect on virion binding to the target cells. Further experiments showed that knockdown of PLSCR1 did not influence CD4 and CXCR4 expression levels on the plasma membrane of SupT1 cells (**Fig. 2C**).

**Figure 2.**
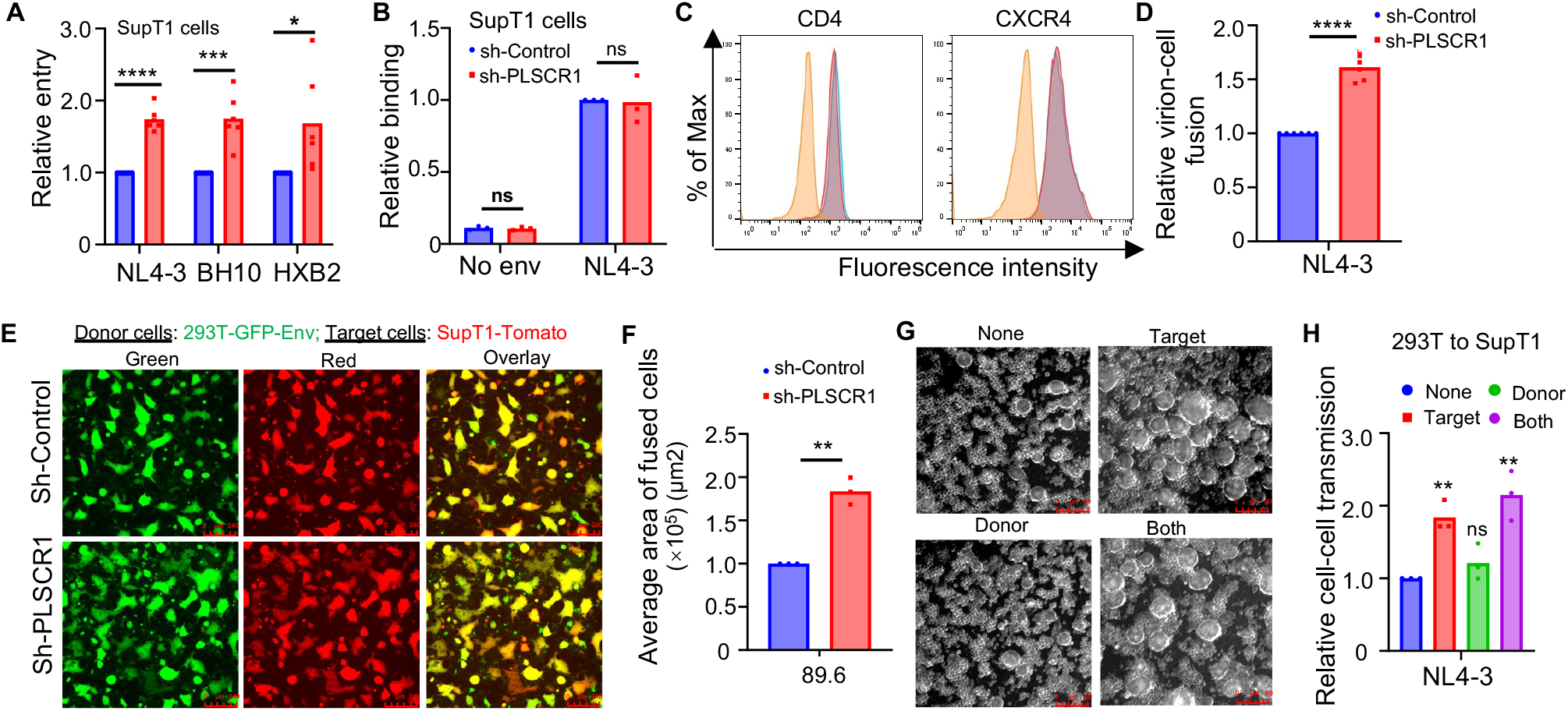
PLSCR1 inhibits HIV-1 entry and cell-to-cell transmission by blocking Env-mediated fusion. **(A)** Entry of HIV-1 pseudoviruses bearing NL4-3, BH10, or HXB2 Env in SupT1 cells with or without PLSCR1 knockdown. **(B)** Binding of GFP-expressing, HIV-1 NL4-3 Env-bearing virus-like particles to SupT1 cells as determined by flow cytomentry. **(C)** The surface level of CD4 and CXCR4 in SupT1 cells stably expressing shControl or shPLSCR1 as assessed by flow cytometry. **(D)** Virion–cell fusion measured by HIV-1 BlaM-Vpr assay in SupT1 cells expressing shControl or shPLSCR1. **(E, F)** Env-mediated cell–cell fusion assessed by co-culturing 293T-GFP (donor) and SupT1-Tomato (target) cells; fusion was imaged and quantified by averaging fused areas. **(G, H)** Knockdown of PLSCR1 in target cells, not in donor cells, enhances Env-mediated cell-to-cell transmission. Representative brightfield images of syncytia formation between 293T donor cells and SupT1 target cells are shown under different PLSCR1 knockdown conditions as follows: donor cells only, target cells only, both donor and target cells, or none (**G**). Quantification of *Gaussia* luciferase activity in co-culture supernatants, which is normalized to the “None” condition **(H)**. Data represent means ± SEM from three or more independent experiments. Statistical significance was determined using unpaired two-tailed Student’s t-test. *p < 0.05, **p < 0.01, ***p < 0.001, ****p < 0.0001. ns, not significant.

We next determined whether PLSCR1 affects HIV-1 Env-mediated membrane fusion. To assess virus–cell fusion, we employed the β-lactamase (BlaM)-Vpr-based HIV-1 fusion assay, in which the BlaM-Vpr chimeric protein is incorporated into HIV-1 virions and subsequently delivered into the cytosol of target cells upon viral fusion. As shown in **Fig. 2D**, knockdown of PLSCR1 in SupT1 cells resulted in a 1.6-fold (p < 0.0001) increase in virion–cell fusion. We further assessed the effect of PLSCR1 on Env-mediated cell-cell fusion by transfecting HEK293T-GFP cells with dual-tropic HIV-1 Env 89.6, followed by co-culturing them with SupT1-tomato cells transiently transduced to express either shRNA-Control or shRNA-PLSCR1. Fusion events were monitored over time microscopically and quantified. We found that PLSCR1 knockdown led to a ∼1.8-fold enhancement of Env-mediated cell-cell fusion (p < 0.01) (**Fig. 2E-F**).

Finally, we evaluated the possible role of PLSCR1 in HIV-1 cell-to-cell transmission using an intron-containing *Gaussia* luciferase (inGluc) HIV-1-based lentiviral reporter system. The reporter carries an intron that is spliced out post-transcriptionally in target cells upon infection, yielding functional luciferase; hence, Gluc is not expressed in donor cells, thus facilitating the measurement of cell-to-cell transmission. 293T donor cells stably expressing either shControl or shPLSCR1 were co-transfected with HIV-inGluc and NL4-3 Env, then co-cultured with SupT1 target cells also stably expressing shControl or shPLSCR1 for 24–36 hours. Gaussia luciferase activity in the culture supernatant was measured to assess cell-to-cell transmission efficiency. We found that PLSCR1 knockdown in SupT1 target cells, but not in 293T donor cells, significantly increased cell-to-cell transmission, as indicated by elevated Gaussia luciferase activity (p < 0.01), accompanied by enhanced cell–cell fusion (**Fig. 2G–H)**. These findings demonstrate that PLSCR1 restricts both HIV-1 cell-to-cell transmission and Env-mediated membrane fusion.

Our study identifies PLSCR1 as an ISG that restricts lentiviral entry and cell-to-cell transmission by targeting the membrane fusion step. These findings provide new insights into the antiviral function and mechanism of PLSCR1, which appears to broadly inhibit viral infection by interfering with conserved features of the fusion process, potentially through modulation of membrane biophysical properties and/or interactions with viral fusion proteins. Further investigation is warranted to elucidate the precise molecular and biophysical mechanisms by which PLSCR1 inhibits viral entry and fusion.

## MATERIALS AND METHODS

### Cells, plasmids and reagents

HEK293T and HeLa-TZM cells were maintained in DMEM supplemented with 0.5% penicillin/streptomycin and 10% fetal bovine serum (FBS). SupT1, SupT1-R5 and SupT1-tomato were maintained in RPMI 1640 supplemented with 0.5% penicillin/streptomycin and 10% FBS. Purified human CD4+ T lymphocytes (Charles River, ID D238769, 278465 and 329252) were activated with phytohemagglutinin of phaseolus vulgaris (Sigma-Aldrich, 5 μg/mL) and 10 U/mL interleukin-2 (R&D Systems, Cat#202-IL-010) for 3 days and then maintained in RPMI1640 with 10 U/mL interleukin-2 till use. SupT1 cells stably expressing shRNA-Control and shRNA-PLSCR1 cell lines were generated by lentiviral transduction using shRNA (Sigma; SHC002 and TRCN0000056271), followed by selection with puromycin (1 μg/mL; Sigma, P8833). SupT1-R5 and SupT1-Tomato cells were transiently transduced with the same lentiviral vectors expressing shRNA-Control or shRNA-PLSCR1 for 24 h before infection. Knockdown efficiency was assessed by western blotting or reverse transcription-quantitative PCR (RT-qPCR) 48 h post-transduction using the following primers: PLSCR1 forward, 5′-CGGCAGCCAGAGAACTGTTTTA; reverse, 5′-CTTGGAATGCTGTCGGTGGA. Human β-actin forward, 5′-GGACTTCGAGCAAGAGATGG; reverse, 5′-AGCACTGTGTTGGCGTACAG.

The proviral HIV-1 pNL4-3 (Cat#114) (16), p89.6 (Cat#3552) (17), pNL(AD8) (Cat#11346) (18), BaL, HIV-2_Rod-1_ (Cat#207), SIVagm (Cat#3444), HIV-1 92/UG/021 (Cat#1648), HIV-1 93/MW/965 (Cat#2913) and HIV-1 93/BR/020 (Cat#2329) were obtained from the BEI/NIH AIDS Reagent Program. 89.6 Env expression vector was kindly provided by Dr. Jesse Kwiek (The Ohio State University, Columbus, OH, USA). The HXB2 Env expression vector was obtained from Dr. Chen Liang’s lab (McGill University, Montreal, QC, Canada). The HIV-1 inGLuc vector was obtained from David Derse (National Cancer Institute, Frederick, MD) (19).

Antibodies against HIV-1 Gag p24 (Cat#1513), CD4 (Cat#724), and CXCR4 (Cat#4083) were obtained from BEI/NIH AIDS Reagents Program. The anti-PLSCR1 antibody was purchased from Proteintech (catalog no. 11582-1-AP), and anti-β-actin antibody from Santa Cruz Biotechnology (Cat#sc-47778). Recombinant human IFN-β was obtained from Alpha Diagnostic Intl. (Cat#48362). FITC-conjugated secondary antibodies against mouse, rabbit, or sheep IgG were from Sigma-Aldrich.

### Transfection and virus production

For infectious virus production, HEK293T cells were transfected with proviral plasmids encoding various strains of HIV-1, HIV-2, or SIV using polyethylenimine (PEI; Polysciences) according to the manufacturer’s instructions. For pseudotyped virus production, pLenti-puro-GFP-based lentiviral particles were generated by co-transfecting 293T cells with plasmids encoding HIV-1 Env, pLenti-puro-GFP, and HIV-1 Gag-Pol Δ8.2 at a ratio of 0.5:1:1. Lentiviral shRNA particles were produced similarly by transfecting pLenti-shRNA, HIV-1 Gag-Pol Δ8.2, and pMD.G at a 1:1:0.5 ratio. Intron-containing Gaussia luciferase (inGluc) reporter virus stocks were produced by transfecting HEK293T cells with pHIV-inGluc, HIV-1 Gag-Pol Δ8.2, and the HIV-1 Env expression vector at a ratio of 1:1:0.5. All virus-containing supernatants were collected at 48 and 72 h post-transfection, clarified by centrifugation, filtered through a 0.45-μm membrane, and stored at -80°C. Virus-like particles (VLPs) for binding assays were generated by co-transfecting 293T cells with HIV-Gag-GFP and NL4-3 Env plasmids at a 1:1 ratio. VLPs were purified by ultracentrifugation at 25,000 rpm for 2 h at 4°C over a 20% sucrose cushion and stored at -80°C until use.

### Infection of CD4^+^ T lymphocytes

Primary human CD4^+^ T lymphocytes were activated for 3 days with 5 μg/mL Phaseolus vulgaris lectin (Sigma-Aldrich) and 10 IU/mL recombinant human interleukin-2 (IL-2; R&D Systems). Cells were then transduced with lentiviral pseudotypes expressing either control shRNA or PLSCR1-targeting shRNA for 24 h, followed by selection with puromycin (1 μg/mL) for an additional 3 days. Cells were subsequently infected with HIV-1 in the presence of 5 μg/mL polybrene (Santa Cruz Biotechnology, catalog no. 134220) by spinoculation at 1,680 × g for 1 h at room temperature. After 6 h of incubation at 37°C, the virus was removed by washing with phosphate-buffered saline (PBS), and cells were resuspended in fresh RPMI 1640 medium supplemented with 10 IU/mL IL-2, which was maintained throughout the experiment.

*Long-term HIV-1 replication in SupT1 cells*

SupT1 shRNA-Control and SupT1 shRNA-PLSCR1 cells were infected with NL4.3 equivalent to 5 ng of viral p24 by spinoculation. Six hours after infection, unbound virus was removed by washing with PBS. Virus particles secreted into the culture medium were collected every 2 days for a period of 14 days. Harvested cell supernatants were centrifuged at 3500 rpm for 10 min to remove the cell debris, and viral p24 was measured by p24 ELISA or infectivity was determined by infecting HeLa-TZM cells, respectively.

### Viral infectivity assay

HIV-1 infectivity was determined by infecting indicator HeLa-TZM cells. Briefly, 36 – 48 h after infection, cells were washed once with PBS and lysed with lysis buffer (50 mM Tris (pH 7.4), 150 mM NaCl, 1 mM EDTA, 1 mM EGTA, and 1% Triton X-100). Approximately 20 ul of cell lysates was incubated with 20 ul of firefly luciferase substrates to determine HIV-1 infectivity.

### Western blotting

Virus-infected cells were lysed in prechilled radioimmune precipitation assay buffer (1% Nonidet P-40, 50 mM Tris-HCl, 150 mM NaCl, 0.1% SDS, and protease inhibitor mixture) for 40 min, and protein samples were subjected to 10% SDS-PAGE. Proteins were transferred to a PVDF membrane, and detected by using anti-HIV-1 p24, anti-PLSCR1, or anti-β-actin antibodies, and protein signals were detected using a Fuji Film imaging system.

### ELISA

The concentration of HIV-1 p24 antigen in culture supernatants was measured using the HIV-1 p24 ELISA Pair Set (Sino Biological Inc., Cat# SEK11695), following the manufacturer’s instructions. Briefly, 96-well plates were coated with capture antibody overnight at 4°C, blocked, and incubated with appropriately diluted supernatants or p24 standards. After washing, detection antibody and streptavidin-HRP were sequentially added. The signal was developed using TMB substrate and stopped with sulfuric acid, and absorbance was measured at 450 nm using a microplate reader. p24 concentrations were determined by comparison to a standard curve.

### Virus purification

For viral binding and virion–cell fusion assays, VLPs or infectious virus stocks were purified by ultracentrifugation prior to use. Briefly, cell culture supernatants containing VLPs or virus were collected 48 h post-transfection or infection, clarified by low-speed centrifugation (1,500 × g for 10 minutes), and filtered through a 0.45 μm membrane. The clarified supernatant was then layered onto a 20% (w/v) sucrose cushion in PBS and subjected to ultracentrifugation at 25,000 × g for 2 h at 4°C using a SW28 Ti rotor (Beckman Coulter). Pelleted virions were resuspended in cold PBS, aliquoted, and stored at –80°C until further use. Virus concentration was quantified by p24 ELISA prior to downstream applications.

### Viral entry and binding assay

For viral entry assay, SupT1 cells expressing shRNA control or PLSCR1 were infected with pLenti-GFP pseduvirions bearing different HIV-1 Env for 48 h. For virion binding assay, SupT1 cells expressing shRNA control or PLSCR1 were incubated with purified HIV-1 Gag-GFP based VLPs at 4°C for 2 h. After three washes with cold PBS to remove unbound virus, cells were fixed with 3.7% formaldehyde for 10 min. For both assays, cells were subjected to flow cytometric analysis.

### Cell surface staining

To assess the surface expression of CD4 and coreceptor CXCR4, SupT1 cells were harvested, washed with ice-cold PBS containing 2% FBS, and incubated with fluorophore-conjugated antibodies against human CD4 (Cat#724) or CXCR4 (Cat#4083) for 1 h at 4°C in the dark. After staining, cells were washed twice with FACS buffer (PBS + 2% FBS) and analyzed on a flow cytometer. Data were processed using FlowJo software. Appropriate isotype controls were included to define background fluorescence.

### Virion-cell fusion

The β-lactamase (BlaM)-Vpr-based HIV-1 virion-cell fusion assay was performed as previously described (20). Briefly, HIV-1 particles were produced by co-transfecting 293T cells with HIV-1 NL4-3 and a BlaM-Vpr expression construct (Gift of Warner Greene). Viral supernatants were collected 48 h and 72 h post-transfection, purified by ultracentrifugation through a sucrose cushion and quantified by p24 ELISA. Target cells incubated with BlaM-Vpr-containing HIV NL4-3 virus for 2 h at 37°C. Following infection, cells were washed and loaded with the CCF2-AM substrate (Thermo Fisher Scientific, Cat#K1085) according to the manufacturer’s instructions. Cells were then incubated with the substrate for 2 h at room temperature in the dark. The substrate was subsequently removed, and cells were resuspended in CO_2_-independent medium supplemented with 10% FBS and 0.5 mM probenecid. Cells were further incubated for 12–16 h at room temperature in the dark. Finally, cells were fixed and analyzed by flow cytometry. Fusion efficiency was quantified as the percentage of cells exhibiting a fluorescence emission shift from green (520 nm) to blue (447 nm), indicative of CCF2 cleavage by BlaM delivered upon viral fusion.

### Cell-cell fusion

To assess HIV-1 Env-mediated cell-cell fusion, 293T-GFP cells were transfected with HIV-1 Env (strain 89.6) expression vector or an empty vector as a negative control. At 24 h post-transfection, donor 293T-GFP cells were harvested and co-cultured with SupT1-Tomato cells that had been transiently transfected with either control shRNA or shRNA targeting PLSCR1. Co-cultures were incubated at 37°C for 4–6 h to allow membrane fusion. Fusion events were assessed by fluorescence microscopy. Syncytium formation was quantified by measuring the area of overlapping GFP^+^/Tomato^+^ cells using ImageJ software. The extent of fusion was expressed as the total fused area per field or as fold change relative to control conditions.

### Cell-to-cell transmission

Cell-to-cell transmission of HIV-1 was evaluated using an intron-containing Gaussia luciferase (inGluc) HIV-1 reporter system (21). Briefly, donor 293T cells expressing shRNA-Control or PLSCR1 were co-transfected with the HIV-inGluc reporter plasmid and HIV-1 Env expression vector (NL4-3 Env). At 24 h post-transfection, donor cells were digested with PBS containing 5 mM EDTA and co-cultured with SupT1 cells stably expressing either control shRNA or PLSCR1-targeting shRNA. Co-cultures were incubated for 48 h at 37°C, after which supernatants were collected and assayed for *Gaussia* luciferase activity using a luciferase assay system. Luminescence was measured using a plate reader, and transmission efficiency was quantified based on relative light units (RLU).

### Statistical analysis

All statistical analyses were carried out in GraphPad Prism9, with unpaired two-tailed Student’s t test used unless otherwise noted. Typically, data from three to six independent experiments were used for the analysis.

